# Pharmacological characterization of the endocannabinoid sensor GRAB_eCB2.0_

**DOI:** 10.1101/2023.03.03.531053

**Authors:** Simar Singh, Dennis Sarroza, Anthony English, Maya McGrory, Ao Dong, Larry Zweifel, Benjamin B. Land, Yulong Li, Michael R. Bruchas, Nephi Stella

## Abstract

**Introduction:** The endocannabinoids (**eCBs**), 2-arachidonoylglycerol (**2-AG**) and arachidonoyl ethanolamine (**AEA**), are produced by separate enzymatic pathways, activate cannabinoid receptors with distinct pharmacology, and differentially regulate pathophysiological processes. The genetically encoded sensor, GRAB_eCB2.0_, detects real-time changes in eCB levels in cells in culture and preclinical model systems; however, its activation by eCB analogues produced by cells and by phyto-cannabinoids remains uncharacterized, a current limitation when interpreting changes in its response. This information could provide additional utility for the tool in in vivo pharmacology studies of phyto-cannabinoid action.

**Methods:** GRAB_eCB2.0_ was expressed in cultured HEK293 cells. Live cell confocal microscopy and high-throughput fluorescent signal measurements.

**Results:** 2-AG increased GRAB_eCB2.0_ fluorescent signal (EC_50_ = 85 nM), and the cannabinoid 1 receptor (**CB_1_R**) antagonist, SR141617, decreased GRAB_eCB2.0_ signal (**SR1**, IC_50_ = 3.3 nM), responses that mirror their known potencies at cannabinoid 1 receptors (**CB_1_R**). GRAB_eCB2.0_ fluorescent signal also increased in response to AEA (EC_50_ = 815 nM), the eCB analogues 2-linoleoylglycerol and 2-oleoylglycerol (**2-LG** and **2-OG**, EC_50_s = 1.5 and 1.0 μM, respectively), Δ^9^-tetrahydrocannabinol (**Δ^9^-THC**) and **Δ^8^-THC** (EC_50_s = 1.6 and 2.0 μM, respectively), and the artificial CB_1_R agonist, CP55,940 (**CP**, EC_50_ = 82 nM); however their potencies were less than what has been described at CB_1_R. Cannabidiol (**CBD**) did not affect basal GRAB_eCB2.0_ fluorescent signal and yet reduced the 2-AG stimulated GRAB_eCB2.0_ responses (IC_50_ = 8.8 nM).

**Conclusions:** 2-AG and SR1 modulate the GRAB_eCB2.0_ fluorescent signal with EC_50_s that mirror their potencies at CB_1_R whereas AEA, eCB analogues, THC and CP increase GRAB_eCB2.0_ fluorescent signal with EC_50_s significantly lower than their potencies at CB_1_R. CBD reduces the 2-AG response without affecting basal signal, suggesting that GRAB_eCB2.0_ retains the negative allosteric modulator (**NAM**) property of CBD at CB_1_R. This study describes the pharmacological profile of GRAB_eCB2.0_ to improve interpretation of changes in fluorescent signal in response to a series of known eCBs and CB_1_R ligands.

## Introduction

Many physiological functions and behaviors are differentially controlled by endogenously produced AEA and 2-AG that activate CB_1_R.^1^ Specifically, AEA or 2-AG production by select cells will partially or fully activate CB_1_Rs (EC_50_s: ≈10-100 nM and 30-300 nM, respectively) in an autocrine and paracrine fashion.^2^ The effects of CB_1_R activation are dependent on both cell type and coupling to intracellular signaling systems: CB_1_R are expressed at remarkably different levels by distinct excitatory and inhibitory neurons and by glial cells where they couple to specific signaling pathways. Thus, cell specific differential activation of CB_1_Rs by AEA and 2-AG in brain fine-tunes excitatory and inhibitory neurotransmission and neuromodulation, and regulates neuronal metabolism and phenotype.^3;4^ This fundamental signaling mechanism is modulated by Δ^9^-THC through its binding to the orthosteric binding site of CB_1_R where it acts as a partial agonist, and by cannabidiol (**CBD**) that interacts with a putative allosteric binding site on CB_1_R.^5^ For example, free CB_1_Rs expressed throughout the brain are partially activated by THC whereas localized activity-dependent increases in 2-AG and stimulation of CB_1_R signaling is reduced by THC.^6;7^ Multiple lines of evidence suggest that CBD acts as a NAM of CB_1_R, though direct CBD binding to CB_1_R has still not been demonstrated.^8;9^ Thus, CBD reduces 2-AG-stimulated CB_1_R signaling without influencing basal/tonic CB_1_R signaling. This premise emphasizes a need to better understand the role of endogenously produced AEA and 2-AG, their dynamics, and how the presence of THC and CBD affects their activation of the CB_1_R.

Mass spectrometry has demonstrated that 2-AG is 10-1000 times more abundant than AEA in select cell types and tissues; however, little is known about their activity-dependent and spatio-temporal changes that occur within seconds.^1;10^ Importantly, increased cellular activity not only enhances the production of AEA and 2-AG, but also enhances the production of lipid analogues synthesized by the same enzymatic pathways. For example, 2-LG and 2-OG activate CB_1_R yet with lower potency and efficacy than 2-AG.^11–13^ Thus, localized activity-dependent increases in eCBs and their analogues will differentially activate CB_1_R within seconds. The recent development of genetically encoded fluorescent sensors has enabled the real-time detection of changes in the levels of neurotransmitters and neuromodulators in live tissues.^14^ This technology leverages the selective binding of endogenous agonists to specific receptors that stabilize their conformation. For example, the GRAB_eCB2.0_ sensor was recently engineered starting from CB_1_R by introducing a circularly permutated-green fluorescent protein (**cpGFP**) in its third intracellular loop.^15;16^ Thus, GRAB_eCB2.0_ was developed by screening for constructs with functional insertion sites of cpGFP, followed by individual randomized mutations of amino acids that increased the fluorescent signal in response to 2-AG specifically.^17^ Several laboratories reported that GRAB_eCB2.0_ signal increases within seconds when exogenously applying 2-AG or AEA to cells in culture, as well as when endogenously stimulating eCB production in cells in culture, mouse brain slices and behaving animals.^15;18–20^

In the current study, we measured GRAB_eCB2.0_ signal in HEK293 cells in culture using live cell fluorescence microscopy and a high-throughput fluorescence plate reader assay. We found that 2-AG and SR1 formulated in buffer containing bovine serum albumin (**BSA**), a lipid binding protein known to assist eCB’s activation of CB_1_R, modulate GRAB_eCB2.0_ fluorescent signal in HEK293 cells with potencies that closely mirror their reported activities at CB_1_R.^15;21^ Thus, we leveraged this experimental approach to characterize the pharmacological profile of eCB analogues and phyto-CBs at GRAB_eCB2.0_.

## Materials and Methods

### Chemicals and Reagents

2-AG (Cayman), 1-arachodonoylglycerol (Cayman), Δ^9^-THC (NIDA Drug Supply Program), CP55940 (Cayman), SR141716 (NIDA Drug Supply Program), arachidonoyl ethanolamine (Cayman), goat anti-CB_1_R antibody (1:1000 for ICC and 1:2,500 for immunoblotting; gift from Dr. Ken Mackie); AlexaFluor 647 conjugated donkey anti-goat (1:1000; Invitrogen); IRDye 800 CW conjugates donkey anti-goat (1:10,000; LI-COR).

### Cloning

GRAB_eCB2.0_ and mut-GRAB_eCB2.0_ DNA were subcloned into an AM/CBA-WPRE-bGH plasmid using the BamHI and EcoRI restriction sites. The plasmid was purified (Purelink HiPure Plasmid Maxiprep Kit, Invitrogen, CA) from transformed Stellar Competent Cells (Takara Bio Inc, Japan). The DNA was sequenced and verified (CLC Sequence Viewer 8) prior to use in transfection.

### Cell Culture

HEK293 cells were grown in DMEM (supplemented with 10% fetal bovine serum and 1% penicillin/streptomycin) at 37°C and 5% CO2. To passage cells for experiments, a confluent 10 cm plate of cells was detached by incubating with 0.25% Trypsin-EDTA for 2-3 min at 37°C, adding 4-5 ml of supplemented DMEM and using gentle pipetting to remove any cells still attached, then added to a new plate with fresh supplemented DMEM. Cells were passaged every 3-4 days, and for no more than 25 passages.

### Transfection

All transfections were performed by incubating DNA with the transfection reagent polyethylenimine (PEI, 25K linear, Polysciences 23966) in a 1:3 ratio in serum free DMEM, incubating for 20-30 min. The DNA/PEI mixture was then added to cells in a dropwise fashion without changing the growth media. Cells were transfected when reaching >50% confluent and were incubated for 24 h after transfection before harvesting or using for GRAB_eCB2.0_ assays.

### Immunocytochemistry (ICC)

Glass coverslips (Fisher Scientific, 12-545-82) in a 6-well plate were coated with poly-D-lysine (50 ng/ml, Sigma, P6407) for 1-2 h at 37°C, after which the coverslips were washed 3X with sterile water and 1X with DMEM. HEK293 cells were detached and resuspended in supplemented DMEM as described above, plated at a density of 100,000 cells/well, and were transfected after 24 h with 0.75 μg DNA. Twenty h after transfection, media was removed, and cells were fixed with 4% paraformaldehyde in PBS (Alfa Aeser) for 20 min at room temperature. Following fixation, cells were washed 5X with PBS and permeabilized and blocked with 0.1% saponin (Sigma-Aldrich) made and 1% BSA in PBS for 30 min at room temperature. Cells were incubated in goat anti-CB_1_R antibody (1:1000) overnight at 4°C. Cells were then washed with PBS 6X and incubated in AlexaFluor647 conjugated donkey anti-goat secondary antibody (Invitrogen, 1:1000) for 1 h at room temperature. Cells were washed with PBS 6X, air dried overnight, and mounted using ProLong Diamond Antifade Mountant with DAPI (ThermoFisher, P36966). Images were captured using a LeicaSP8X confocal microscope and a 40X oil objective.

### Western Blotting

HEK293 cells were plated as described above at a density of 500,000 cells/well in a 6-well plate 24 h after plating and were transfected the following day with 0.75 μg DNA. Twenty h after transfection, cells were washed 3X with ice cold PBS, and in the last wash cells were harvested with a cell scraper and pelleted by centrifuging at 500 x g for 10 min. The supernatant was discarded, and cell pellets kept at −80°C until further use. Thus, pellets were thawed on ice, resuspended in lysis buffer (25 mM HEPES pH 7.4, 1 mM EDTA, 6 mM MgCl2, and 0.5% CHAPS), Dounce homogenized on ice (20-30 strokes), and incubated on a rotator at 4°C for 1 h. Homogenates were then centrifuged at 700 x g for 10 min at 4°C, supernatant collected, and protein concentration of supernatant determined using a DC Protein Assay. Samples were then mixed with 4X Laemmli Sample Buffer containing 10% β-mercaptoethanol and incubated at 65°C for 5 min. Twenty-five μg of protein were loaded onto a 10% polyacrylamide gel and transferred to PVDF membrane. After transfer, membrane was washed once with tris-buffered saline (TBS) and incubated in blocking buffer (5% BSA in TBS) for 1 h at room temperature, followed by incubation in goat anti-CB_1_R antibody (1:1000) overnight at 4°C. After incubation in primary antibody, membranes were washed with TBS with 0.05% TWEEN-20 (TBST) 3 X, 10 min each. Membranes were then incubated in IRDye 800 CW conjugates donkey anti-goat (1:10,000) for 1 h at room temperature. Blots were then washed with TBST 3 X, 10 min each followed by 3 washes with TBS, 10 min each. Fluorescent signal was detected using a Chemidoc MP (Biorad). Primary antibodies were diluted in blocking buffer. Secondary antibodies were diluted in 1:1 TBS:Odyssey blocking buffer (LI-COR, 927-50000).

### Live-cell imaging

Glass bottom cell culture plates (MatTek, P35G-1.5-14-C) were coated with poly-D-lysine (50 ng/ml, Sigma, P6407) for 1-2 h at 37°C, after which the poly-D-lysine was removed, and coverslips were washed 3 X with sterile water and one time with DMEM. HEK293 cells were detached and resuspended in supplemented DMEM as described above, counted using a hemocytometer, plated (250,000 cells per well) and were transfected after 24 h with 0.75 μg DNA. Twenty-four h after transfection, media was replaced by serum free DMEM and cells were incubated at 37°C and 5% CO_2_ for 1-2 h. To image, the serum free DMEM was replaced by phosphate-buffered saline (room temperature). The plates were transferred to the microscope (LeicaSP8X) and cells were imaged using a 40X oil objective with the following settings: 485 excitation and 525 emission wavelength, 5% laser power, HyD hybrid detector and a scan speed of 200 lines Hz (0.388 frames per second) with bidirectional scanning. All agents were prepared in PBS supplemented with 1 mg/ml BSA and spiked into buffer of cell on the microscope stage (Final concentration BSA = 0.1 mg/ml).

### 96-well Plate Reader

Clear-bottom, black 96-well plates (USA Scientific 5665-5087) were coated with poly-D-lysine (50 ng/ml, Sigma, P6407) for 1-2 h at 37°C, and coverslips were washed 3 X with sterile water and 1 X with DMEM. HEK293 cells were detached using trypsin, resuspended in supplemented DMEM as described above, plated (20,000 cells per well) and transfected after 24 h with 0.1 μg DNA and 0.3 μg of PEI in 10 μl of serum free DMEM. Twenty-four h after transfection, media was replaced with PBS (supplemented with BSA 0.1 mg/ml and at room temperature). Cells were incubated at room temperature for 20 min then a 1 min baseline fluorescent signal reading was obtained using a fluorescent plate reader (i.e., 485 emission and 525 emission filter settings with a 515 nm cutoff, and a speed of 1 reading every 20 sec). Immediately after baseline reading, agents in 1 mg/mL BSA and PBS were spiked into buffer in wells. Approximately 2 min after addition of treatment, the plate was reread for 5 min. For pre-treatment with SR1, this antagonist was prepped in PBS supplemented with BSA 0.1 mg/ml (room temperature) and spiked into the media.

### Statistical Methods

All GRAB_eCB2.0_ fluorescent signals are expressed as ΔF/F_0_ as calculated by MATLAB for the high-throughput fluorescence assay or FIJI ImageJ for live cell confocal imaging. Data are shown as mean + s.e.m. and significance was determined by running a Two-Way ANOVA with Dunnett’s Multiple Comparison test using GraphPad Prism.

## Results

### Real time activation and antagonism of GRAB_eCB2.0_ fluorescent signal in HEK293 cells in culture: Live cell microscopy

HEK293 cells in culture were transfected with *eCB2.0* DNA plasmid constructs containing the chimeric cytomegalovirus-chicken β-actin promoter to drive expression.^22–26^ GRAB_eCB2.0_ expression was confirmed by western blot using a rabbit polyclonal antibody developed against N-terminal amino acids of CB_1_R that remained unchanged in GRAB_eCB2.0_ (**Figure 1A** and Supplementary Figure S1 for amino acid sequence alignment). Fluorescence confocal microscopy analysis of GRAB_eCB2.0_ expression in fixed HEK293 cells using the CB_1_R antibody showed abundant expression in many cells, consistent with transient transfection approaches (**Figure 1B**).

**Figure 1:**
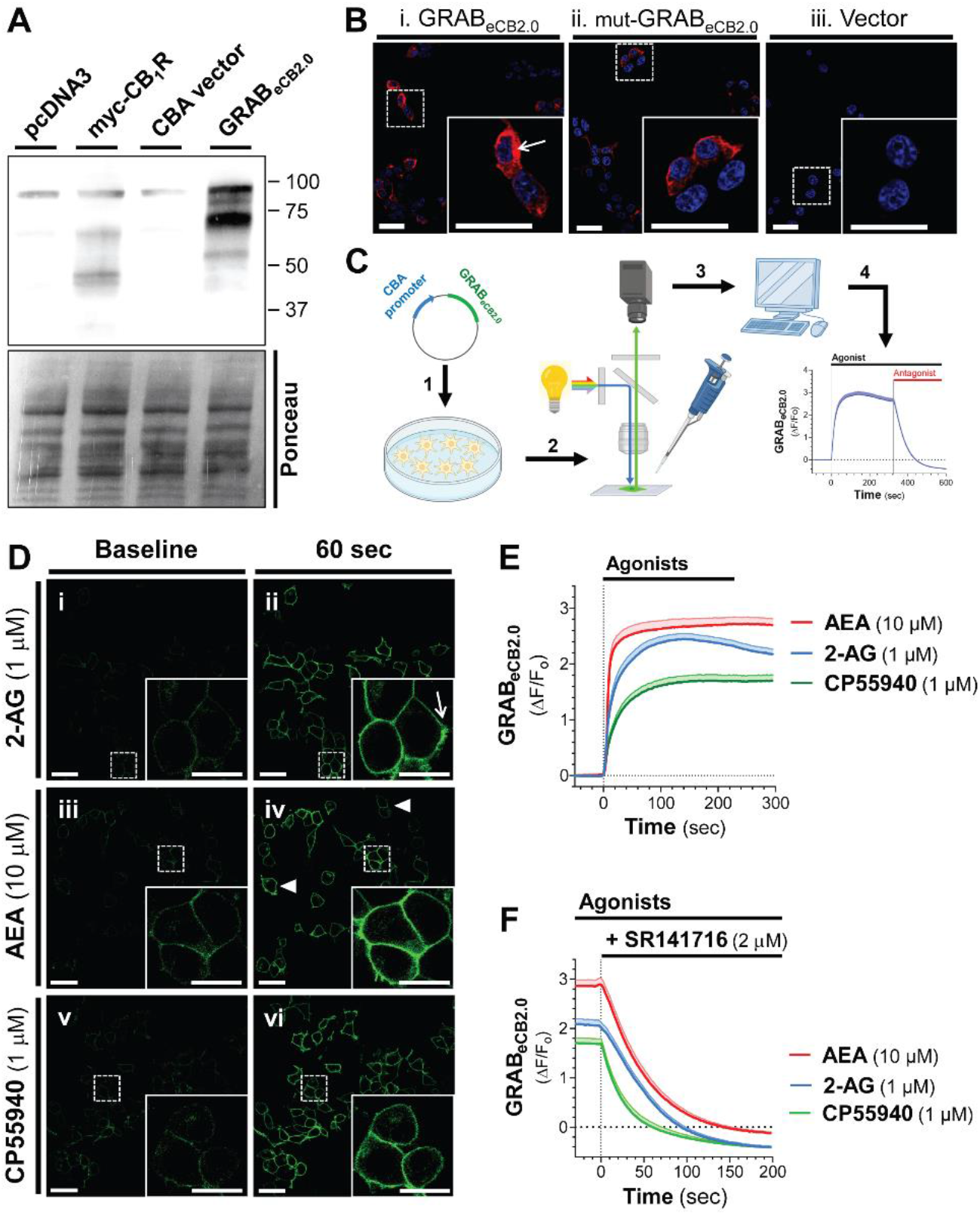
Agonist triggered changes in GRAB_eCB2.0_ fluorescent signal in HEK293 cells detected by livecell confocal microscopy. HEK293 cells were transfected with *eCB2.0* DNA plasmid and GRAB_eCB2.0_ expression was measured by western blot and ICC, and changes in fluorescent signal measured by live-cell confocal microscopy when treated with the CB_1_R agonists 2-arachidonoyl glycerol (**2-AG**), arachidonoyl ethanolamine (**AEA**), or CP55940 (**CP**). **A]** Detection of GRAB_eCB2.0_ expression by western blot using lysates from HEK293 cells transfected with pcDNA3, myc-CB_1_R (positive control), CBA plasmid, or GRAB_eCB2.0_. Both CB_1_R and GRAB_eCB2.0_ were detected using an antibody against the CB_1_R. Loading control: Ponceau stain. **B]** Detection of GRAB_eCB2.0_ (i) or mut-GRAB_eCB2.0_ (ii) expression by ICC of fixed and permeabilized HEK293 cells transfected with *eCB2.0* DNA plasmid (i), mut-*eCB2.0* DNA plasmid (ii), or CBA plasmid (iii). Scale bar = 20 μm (inset scale bar = 20 μm). **C]** Schematic of live cell confocal imaging of HEK293 cells: cells were transfected with *eCB2.0* DNA plasmid (1); 24 h later, cell growth media was exchanged for imaging PBS buffer and cells were placed on a confocal microscope (2); changes in GRAB_eCB2.0_ fluorescent signal was determined by measuring fluorescent signal during baseline, spiking in treatments formulated with BSA (0.1%, 10X), and lastly spiking in SR141716 (**SR1**; 3 and 4). **D]** Live cell images of GRAB_eCB2.0_-expressing HEK293 cells during baseline recording (i, iii, and v) and after 60 sec of treatment with 2-AG (1 μM, ii), AEA (10 μM, iv), or CP (1 μM, vi). Arrows indicate cells with different levels of fluorescent signal to show heterogeneous expression of GRAB_eCB2.0_. Scale bars = 20 μm. **E]** Time courses of GRAB_eCB2.0_ activation (ΔF/F_0_) following treatment with 2-AG (1 μM), AEA (10 μM), and CP (1 μM) as measured by live-cell confocal microscopy. **F]** Effect of SR1 (2 μM) on agonist-stimulated increase in GRAB_eCB2.0_ fluorescent signal. Shaded area represents s.e.m. n= 42-66 cells from 3-5 independent experiments.

We used live-cell confocal microscopy (line scanning frequency: 200 Hz) to establish the time-course of changes in GRAB_eCB2.0_ fluorescent signal in response to 2-AG (the ligand used to develop this sensor) by spiking it directly into the BSA-containing PBS buffer inside the imaging chamber (**Figure 1C**). **Figure 1D_i-ii_** show that HEK293 cells exhibited low, yet clearly detectable basal fluorescent signal at the plasma membrane (treated with vehicle control, DMSO 0.1%); and that 2-AG (1 μM, 60 sec) increased this fluorescent signal. Of note, although the GRAB_eCB2.0_ protein is also expressed in the intracellular compartment of HEK293 cells (see arrow in **Figure 1B** insert), 2-AG only increases GRAB_eCB2.0_ fluorescent signal at the plasma membrane (see arrow in **Figure 1D_ii_** insert). Similarly, AEA (10 μM, 60 sec) and CP (1 μM, 60 sec) increased GRAB_eCB2.0_ fluorescent signal at the plasma membrane (**Figure 1D_iii-iv_**). Basal fluorescent signal and all agonist-triggered increases in GRAB_eCB2.0_ fluorescent signal reached different levels in HEK293 cells as expected by heterologous expression (for example, see **Figure 1D_iv_** arrowheads).

Analysis of the time course of GRAB_eCB2.0_ fluorescent signal (ΔF/F_0_) increase triggered by these agonists indicate that each agonist increased GRAB_eCB2.0_ fluorescent signal within seconds and that these responses plateaued after approximately 60 sec (**Figure 1E**). **Figure 1F** shows that the CB_1_R antagonist, SR1 (2 μM), reduced these responses within 100 sec, and this reduction reached levels below basal. Calculation of the onset of activation (i.e., slope of the fluorescent signal increase within the first 20 sec after start of treatment), the magnitude of the response (i.e., peak fluorescent signal and area under the curve), and the decay following SR1 treatment (i.e., τ value following start of SR1 treatment) indicated that each agonist had significantly different pharmacological profiles at GRAB_eCB2.0_ (**Table 1**). Specifically, 1] AEA triggered a 1.7- and 2.2-fold faster initial response compared to 2-AG and CP respectively, 2] 2-AG and AEA reached similar peak responses, and these were 47-58% greater than the CP maximal response, 3] 2-AG and AEA triggered a 1.4-1.6-fold overall greater response (area under the curve) compared to CP and 4] SR1 antagonized the CP and AEA responses with faster decay than the 2-AG response (24s < 37s < 43s, respectively). Together, these results indicate that GRAB_eCB2.0_ expressed by HEK293 cells are activated by the CB_1_R agonists 2-AG ≈ AEA > CP, and these responses are differentially antagonized by SR1.

**Table 1:** Parameters of GRAB_eCB2.0_ activation and antagonism. Live cell images of GRAB_eCB2.0_-expressing HEK293 cells treated with 2-AG (1 μM), AEA (10 μM), or CP (1 μM), followed by SR1 (2 μM) on agonist-stimulated increase in GRAB_eCB2.0_ fluorescent signal. Calculation of the activation onset (i.e., the slope of the fluorescent signal increase within the first 20 sec after start of agonist treatment), the magnitude of the response (i.e., peak response and area under the curve), and the decay following SR1 treatment (i.e., τ value following treatment). Analysis of data presented in Figure 1, n= 42-66 cells from 3-5 independent experiments.

|  |  | CB<sub>1</sub>R agonists |  |  |
| --- | --- | --- | --- | --- |
| PD parameter | Units | 2-AG | CP | AEA |
| Initial response: slope | ($\times 10^{-2}$ DF/F<sub>0</sub>/sec) | 6.6 | 4.5 | 10.9 |
| Maximal response: peak | (DF/F<sub>0</sub>) | 2.5 | 1.7 | 2.7 |
| Overall response | (area under the curve) | 649 | 453 | 760 |
| Antagonism response: $t$ | (sec) | 42.7 | 24.3 | 36.8 |

### High-throughput measures of GRAB_eCB2.0_ fluorescent signal in HEK293 cells in culture

To further define the pharmacological profile and dynamics of changes in GRAB_eCB2.0_ fluorescent signal, we developed and validated a fluorescent plate reader assay (96 wells, 3 Hz scanning frequency). **Figure 2A** outlines the experimental approach that consists of: 1] transfecting HEK293 cells in 96 well plates with *eCB2.0* DNA plasmid constructs, 2] 24 h after transfection, replacing the cell culture media with buffer (20 min), inserting the plate into the reader and measuring GRAB_eCB2.0_ fluorescent signal for the last 2 min of this preincubation period (i.e., Basal fluorescent signal), 3] promptly removing the plate from the reader, spiking agents (10X concentration) prepared in PBS supplemented with BSA (1 mg/ml) in media, and reinserting the plate in the reader, and finally 4] measuring fluorescence for 5 min (i.e., Stimulation fluorescent signal). To facilitate a data analysis pipeline, we developed a MATLAB R2021a algorithm that averages the fluorescent signal value of each well over time, for multiple experiments and at select timepoints (see the following link for the code: https://github.com/StellaLab/StellaLab.git). **Figures 2B-C** show that 2-AG and AEA induced concentration-dependent increases in GRAB_eCB2.0_ fluorescent signals that were detected at the 0 sec timepoint (i.e., when the first Stimulation fluorescent signal was measured). For example, both the 10 nM and 100 nM 2-AG responses at 0 sec reached a GRAB_eCB2.0_ fluorescent signals value of 0.19 and 0.96 ΔF/F_0_ over basal, respectively; and both these responses reached an initial inflection point at 0.66 and 0.33 min, respectively (**Figures 2B**, see color coded arrows). Of note, 2-AG at 3 μM induced the strongest response that reached an inflection point at ≈0.66 min, whereas 2-AG at 10 μM induced a response had already reached a maximum plateau response at the 0 min timepoint (**Figures 2B**). Thus, 2-AG rapidly activated GRAB_eCB2.0_ in a concentration dependent manner as calculated by its initial response (i.e., slope: ΔΔF/F_0_ between 0 min and the initial inflection time point) (**Figures 2D**). To calculate EC_50_ values, we averaged the GRAB_eCB2.0_ fluorescent signals between 4-5 min, and found 85 nM for 2-AG, a value that is consistent with its reported potency at CB_1_R (e.g., EC_50_ = 12-100 nM) (**Figures 2E**).^27–30^ AEA also induced a concentration-dependent and rapid increase in GRAB_eCB2.0_ fluorescent signal that was detected at the 0 sec timepoint, but only reached a significant initial response starting at 30 nM, and a maximum plateau response at 10 μM (**Figures 2C-D**). Thus, AEA induced a concentration dependent increase in GRAB_eCB2.0_ with an EC_50_ = 815 nM, an activity that is approximately 10-fold less potent than AEA’s potency at CB_1_R (e.g., EC_50_ = 69-276 nM) (**Figures 2E**).^30–32^ Of note, the EC_50_s values of 2-AG and AEA for GRAB_eCB2.0_ measured here are lower than their previously reported values in cell culture models systems (i.e., 2-AG = 3.1-9.0 μM and AEA = 0.3-0.8 μM), most likely because we included BSA in the buffer to facilitate solubility and interaction with GRAB_eCB2.0_.^15;33^

**Figure 2:**
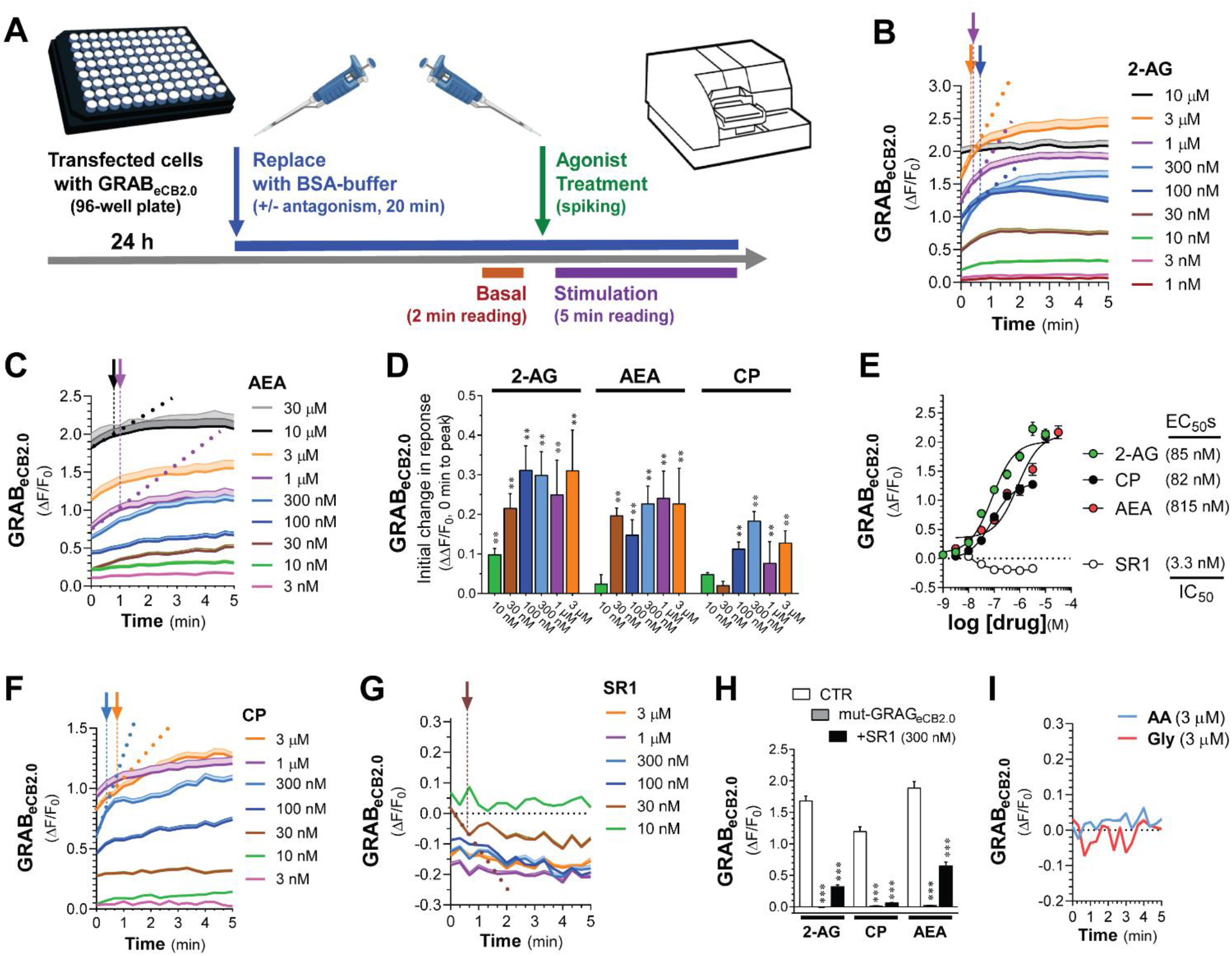
2-AG, AEA, and CP differentially activate GRAB_eCB2.0_: High-throughput fluorescence assay. **A]** matic of high-throughput fluorescent plate reader assay: HEK293 cells were plated in a 96-well plate and transfected with *eCB2.0* DNA plasmid; 24 h post transfection, growth media was replaced with imaging PBS buffer and incubated for 20 min. For the last 2 min of this incubation, the plate was placed in the fluorescent plate reader and a basal fluorescent signal was measured. Treatments were spiked in immediately after the basal reading and the pate was reinserted in the plate reader and fluorescent signal was recorded for 5 min. **B, C, F, G** and **I]** Kinetics of GRAB_eCB2.0_ activation (ΔF/F_0_) following treatment with increasing concentrations of 2-AG (B), AEA (C), CP (F), SR1 (G) and arachidonic acid (**AA**) and glycerol (**Gly**) (I). Arrows represent inflection points and dotted lines represent initial change in response (slope between t=0 and time at inflection). **D]** Initial change in ΔF/F_0_ response elicited by increasing concentration of 2-AG, AEA, or CP shown in (B, C, and F). **E]** Concentration dependent responses and EC_50_s of 2-AG, AEA, and CP at inducing GRAB_eCB2.0_ fluorescent signal and of SR1 at reducing GRAB_eCB2.0_ fluorescent signal as determined by averaging ΔF/F_0_ between 4-5 min. **H]** 2-AG (1 μM), CP (1 μM), and AEA (10 μM)-induced increases in GRAB_eCB2.0_ fluorescent signal was reduced by SR1 (300 nM) and absent in HEK293 cells expressing mut-GRAB_eCB2.0_. Data represents mean ΔF/F_0_ between 4-5 min. Shaded areas on time-course plots and error bars on histograms represent s.e.m. Statistics: (D) **P<0.01 significantly different from basal (Two-Way ANOVA followed by Dunnett’s). (H) ***P<0.001 significantly different from corresponding CTR treatment (Two-way ANOVA followed by Dunnett’s). n= 9-50 independent experiments performed in triplicate.

To further characterize these high-throughput measures, we tested the effect of CP and SR1 on GRAB_eCB2.0_ fluorescent signal. **Figures 2F** shows that CP induced a concentration-dependent increase in GRAB_eCB2.0_ fluorescent signal that was detected at 0 sec and reached an initial inflection time point within ≈1.33 min and an EC_50_ = 82 nM, an activity that is at least 3-fold less potent than CP’s potency at CB_1_R (e.g., EC_50_ = 0.05-31 nM) (**Figures 2D-E**).^34^ As expected, SR1 induced a rapid and concentration-dependent decrease in GRAB_eCB2.0_ fluorescent signal that was detected at 0 sec and plateaued below basal levels starting at 30 nM and for at least 5 min (**Figures 2G**). The IC_50_ for SR1 measured between 4-5 min was 3.3 nM, a value that mirrors SR1’s potency at CB_1_R (IC_50_ = 0.3-17 nM) (**Figures 2E**).^34–37^ Pre-treatment of HEK293 cells with SR1 (300 nM, 20 min) followed by treatment with 2-AG, AEA and CP significantly reduced GRAB_eCB2.0_ activation (**Figures 2H**). **Figure 2H** also shows that 2-AG, AEA and CP failed to elicit increases in fluorescent signals in HEK293 cell expressing the mut-GRAB_eCB2.0_, which has a similar expression profile as GRAB_eCB2.0_ as determined by fluorescence confocal microscopy (**Figure 1B**). Thus, mut-GRAB_eCB2.0_, which has a phenylalanine 177 to alanine mutation in the region within the orthosteric binding pocket to impair ligand binding, represents a valid negative control.^38;39^ As previously reported, the products of 2-AG hydrolysis, arachidonic acid, and glycerol, did not influence the GRAB_eCB2.0_ fluorescent signal (**Figure 2I**).^15^ Together, these results provide strong support for use of this high throughput experimental approach to study the pharmacology and dynamics of changes in GRAB_eCB2.0_ fluorescence when expressed by cells in culture, and show that: 1] Pronounced increases and decreases in GRAB_eCB2.0_ fluorescent signal are reproducibly detected using a 96 well plate-reader format and promptly spiking agents in media, 2] 2-AG and SR1 modulate GRAB_eCB2.0_ fluorescent signal with EC_50_s comparable to their activities at the CB_1_R, and 3] AEA and CP also increase GRAB_eCB2.0_ fluorescent signal, although with 2-10-fold lesser activities than at CB_1_R.

### Activity of 2-AG analogues at GRAB_eCB2.0_

We leveraged the high-thought approach to determine whether 2-LG and 2-OG, which are produced by cells concomitantly to 2-AG, change GRAB_eCB2.0_ fluorescent signals when expressed in HEK293 cells.^11;12^ Specifically, although 2-LG (18:2) and 2-OG (18:1) are lipid analogues of 2-AG (20:4) and are produced by the same biosynthetic and metabolic pathways as 2-AG, they are likely to activate CB_1_R signaling with much lower potency and efficacy than 2-AG. For example, 2-LG partially activates CB_1_R (EC_50_s = 16.6 μM) or antagonizes CB_1_R depending on the model system, while there is still no evidence that 2-OG might also activate CB_1_R.^11;12^ **Figures 3A-B** show that both 2-LG and 2-OG induced a concentration-dependent increase in GRAB_eCB2.0_ fluorescent signal that was detected at 0 sec and that these responses rapidly reached their initial inflection time points within 1 min. Thus, 2-LG and 2-OG activated GRAB_eCB2.0_ with increasingly rapid onset and EC_50_s = 1.0 and 1.5 μM, respectively, activities that are greater than their known potencies at CB_1_R (**Figures 3C-D**). We next tested 1-arachidonoylglycerol (**1-AG**, 20:4), which is a product of a non-enzymatic isomerization of 2-AG that is not endogenously produced by mammalian cells, but is commonly used to study the structure activity relationship of monoacylglycerols at CB_1_R as its acyl chain length and saturation match that of 2-AG (EC_50_s = 1-1.9 μM).^29;40;41^ We found that 1-AG induced a concentration-dependent increase in GRAB_eCB2.0_ fluorescent signal that was detected at 0 sec, reached its initial inflection timepoint within 1 min and an EC_50_ = 1.8 μM, an activity that also mirrors its potency at CB_1_R (**Figures 3D-E**). The responses to these three monoacylglycerols were blocked by pretreatment with SR1 (300 nM) and absent in cells expressing the mut-GRAB_eCB2.0_ (**Figure 3F**). Thus, 1-AG, 2-LG, and 2-OG increase GRAB_eCB2.0_ fluorescent signal (rank order of potency: 2-OG > 2-LG > 1-AG; maximum efficacy: 1-AG >> 2-LG > 2-OG). Notably, all three lipid analogues had lower potencies than 2-AG and AEA, indicating that the GRAB_eCB2.0_ has greater sensitivity for these two eCBs compared to similar monoacylglycerol lipids.

**Figure 3:**
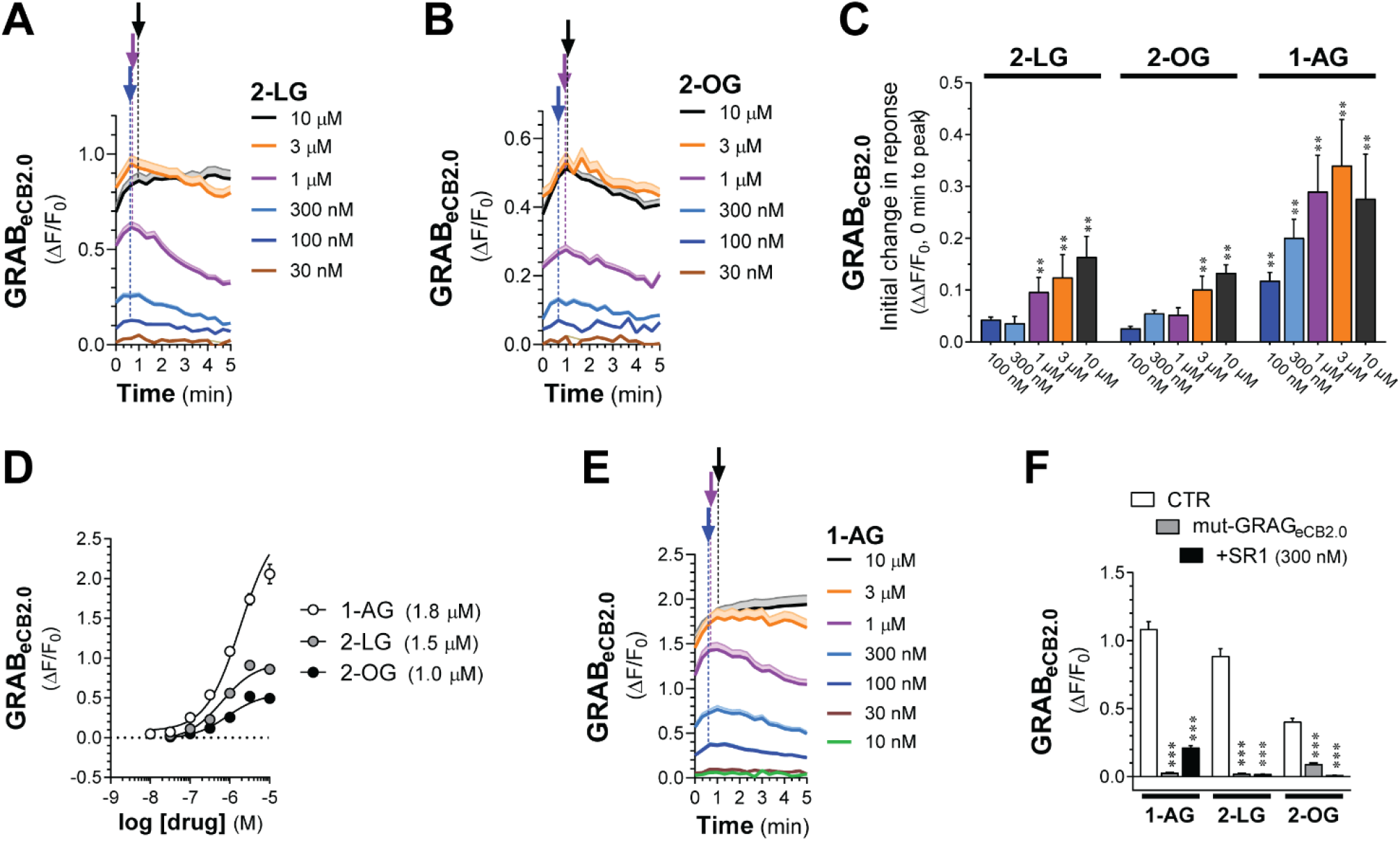
2-LG, 2-OG, and 1-AG activate GRAB_eCB2.0_. HEK293 cells were transfected with *eCB2.0* DNA plasmid and changes in fluorescent signal measured by <u>high-throughput fluorescence assay</u>. **A, B, E]** of GRAB_eCB2.0_ activation (ΔF/F_0_) following treatment with increasing concentrations of 2-LG (A), 2-OG (B), and 1-AG (E). Shaded areas in time courses and error bars represent s.e.m. Arrows represent inflection points and dotted lines represent initial change in response (slope between t=0 and time at inflection). **C]** Initial response (slope between time = 0 and inflection point) of ΔF/F_0_ response. **D]** Concentration dependent responses and EC_50_s of 2-LG, 2-OG, 1-AG at inducing GRAB_eCB2.0_ fluorescent signal as determined by averaging ΔF/F_0_ between 4-5 min. **F]** 2-LG (1 μM), 2-OG (1 μM), and 1-AG (1 μM)-stimulated increases in GRAB_eCB2.0_ fluorescent signal are reduced by SR1 (300 nM) and absent in HEK293 cells expressing mut-GRAB_eCB2.0_. Statistics: (C) **P<0.01 significantly different from basal (Two-way ANOVA followed by Dunnett’s). (D) ***P<0.001 significantly different from corresponding CTR treatment (Two-Way ANOVA followed by Dunnett’s). n= 3-11 independent experiments performed in triplicate.

### THC and CBD activity at GRAB_eCB2.0_

To extend the pharmacological characterization of GRAB_eCB2.0_ using the high-throughput assay, we tested Δ^9^-THC, the principal psychoactive ingredient in *Cannabis* that activates CB_1_R as a high-affinity partial agonist,^34^ and Δ^8^-THC, which activates CB_1_R with a comparable pharmacology as Δ^9^-THC but represents a minor product of most *Cannabis* strains and remains poorly characterized.^42^ Δ^9^-THC increased GRAB_eCB2.0_ fluorescent signal in a concentration-dependent manner that was detected at 0 sec starting at 1 and 3 μM, and these responses reached their initial inflection points within 1 min (**Figure 4A**). Of note, higher concentrations of Δ^9^-THC, i.e., 10 and 30 μM, triggered a response that continuously increased for at least 5 min without apparent inflection (**Figure 4A**). **Figure 4B** shows that Δ^8^-THC induced a concentration-dependent increase in GRAB_eCB2.0_ fluorescent signal that was detected at 0 sec starting at 1 μM, and that all higher concentrations continuously increased for at least 5 min without inflection. Δ^9^-THC and Δ^8^-THC activate GRAB_eCB2.0_ with comparable EC_50_s as calculated between 4-5 min (1.6 μM and 2 μM, respectively; **Figure 4C**). These 2 responses were blocked by pretreatment with SR1 (300 nM) and absent in cells expressing the mut-GRAB_eCB2.0_ (**Figure 4D**). Thus, Δ^9^-THC and Δ^8^-THC similarly increase GRAB_eCB2.0_ fluorescent signal, although with 10-100-fold lesser activities than their potencies at CB_1_R, and most of these responses at micromolar concentrations continuously increase for at least 5 min.

**Figure 4:**
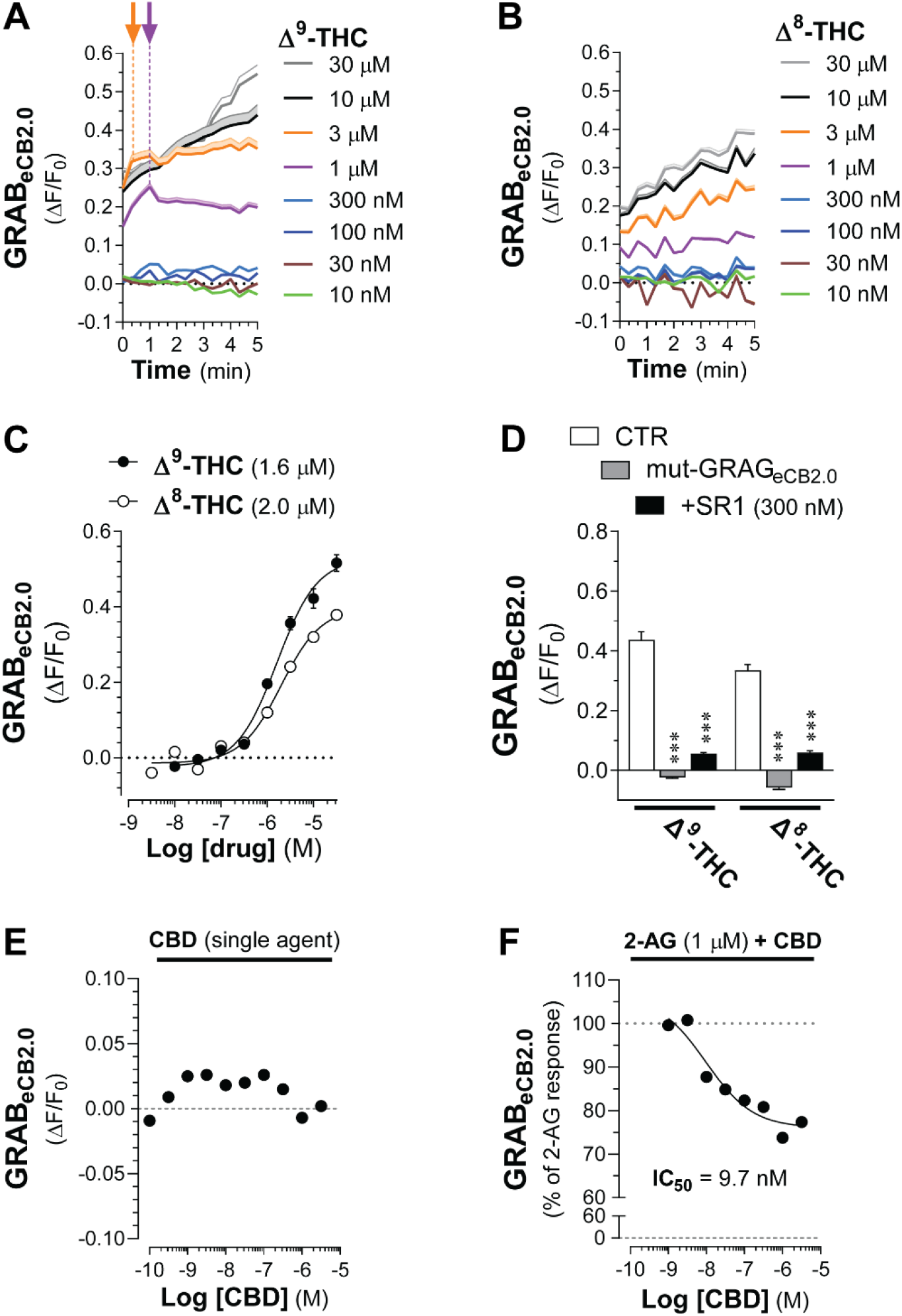
Δ^9^-THC and Δ^8^-THC activate GRAB_eCB2.0_. HEK293 cells were transfected with *eCB2.0* DNA plasmid and changes in fluorescent signal measured by <u>high-throughput fluorescence assay</u>. **A-B]** Kinetics of GRAB_eCB2.0_ activation (ΔF/F_0_) induced by increasing concentrations of Δ^9^-THC (A) or Δ^8^-THC (B). Shaded areas in time courses and error bars represent s.e.m. Arrows represent inflection points and dotted lines represent initial change in response (slope between t=0 and time at inflection). **C]** Concentration dependent responses and EC_50_s of Δ^9^-THC and Δ^8^-THC at inducing GRAB_eCB2.0_ fluorescent signal as determined by averaging ΔF/F_0_ between 4-5 min. d] Δ^9^-THC (10 μM) and Δ^8^-THC (10 μM)-stimulated increases in GRAB_eCB2.0_ fluorescent signal are reduced by SR1 (300 nM) and absent in HEK293 cells expressing mut-GRAB_eCB2.0_. **E-F]** CBD (100 pM-10 μM) does not modulate GRAB_eCB2.0_ fluorescent signal (E) and decreased the 2-AG (1 μM)-stimulated increase in GRAB_eCB2.0_ fluorescent signal (IC_50_ of 9.7 nM). Statistics: (D) ***P<0.001 significantly different from corresponding CTR treatment (Two-way ANOVA followed by Dunnett’s). n= 3-7 independent experiments performed in triplicate.

CBD acts as a NAM of CB_1_R.^8^ We found that increasing concentrations of CBD (from 0.1 nM to 3 μM) did not significantly modulate basal GRAB_eCB2.0_ fluorescent signal (**Figure 4E** and Supplementary Figure S4); however, it significantly reduced the 2-AG-induced increase in GRAB_eCB2.0_ fluorescent signal by 26% at 1 μM and with an IC_50_ = 9.7 nM, values that are similar to CBD’s activity at CB_1_R (**Figure 4E**). Thus, our results suggests that GRAB_eCB2.0_ retains the molecular mechanism that mediates the proposed NAM activity of CBD at CB_1_R.

### Comparing changes in GRAB_eCB2.0_ fluorescent signals

We sought to compare key pharmacological and dynamic parameters that characterize changes in GRAB_eCB2.0_ fluorescent signal elicited by the agents tested in this study (chemical structures in Supplementary S5). **Figure 5A** shows GRAB_eCB2.0_ fluorescent signal increased by each agonist applied at a concentration that approached their EC_50_s and the differences in their overall dynamics. Specifically, when analyzing the ΔΔF/F_0_ of these responses between 3-5 min, we found that the responses induced by monoacylglycerols were decaying between 3-5 min and thus had reached an earlier maximum response, whereas the GRAB_eCB2.0_ fluorescent signals induced by AEA, CP and Δ^9^-THC continuously increased between 3-5 min and thus had not reached a maximum response within this time period (**Figure 5B**). Analysis of dose-responses indicated that 2-AG at 3 μM resulted in the most pronounced increase in GRAB_eCB2.0_ fluorescent signal as measured by area under the curve between 0-5 min, and thus we calculated the relative efficacy of each agent at the of their maximal response concentration compared to the 2-AG response. **Figure 5C** shows that the maximal GRAB_eCB2.0_ responses to AEA (30 μM), 1-AG (10 μM) and CP (3 μM) reached 91%, 81% and 71% of the 2-AG response, respectively. **Figure 5C** also shows that the maximal GRAB_eCB2.0_ responses to 2-LG (10 μM) and 2-OG (10 μM) only reached 37% and 17% of the 2-AG response, respectively, and that Δ^9^-THC (30 μM) and Δ^8^-THC (30 μM) only reached 22% and 16% of the 2-AG response, respectively. Finally, we compared the antagonism of SR1 (100 and 300 nM) for each ligand. **Figure 5D-E** show that the 2-AG (1 μM), CP (1 μM), 2-LG (10 μM), as well as Δ^9^-THC (10 μM) and Δ^8^-THC (10 μM) we partial antagonized by SR1 100 nM and antagonized by more than 80% with SR1 300 nM. By contrast, 1-AG (10 μM) and 2-OG (10 μM) were strongly antagonized by SR1 at both 100 and 300 nM, and AEA (10 μM) was antagonized by only 65% by both 100 and 300 nM SR1. These results reveal differences in key pharmacological and dynamic parameters that describe changes in GRAB_eCB2.0_ fluorescent signal elicited by eCBs and CB_1_R ligands.

**Figure 5:**
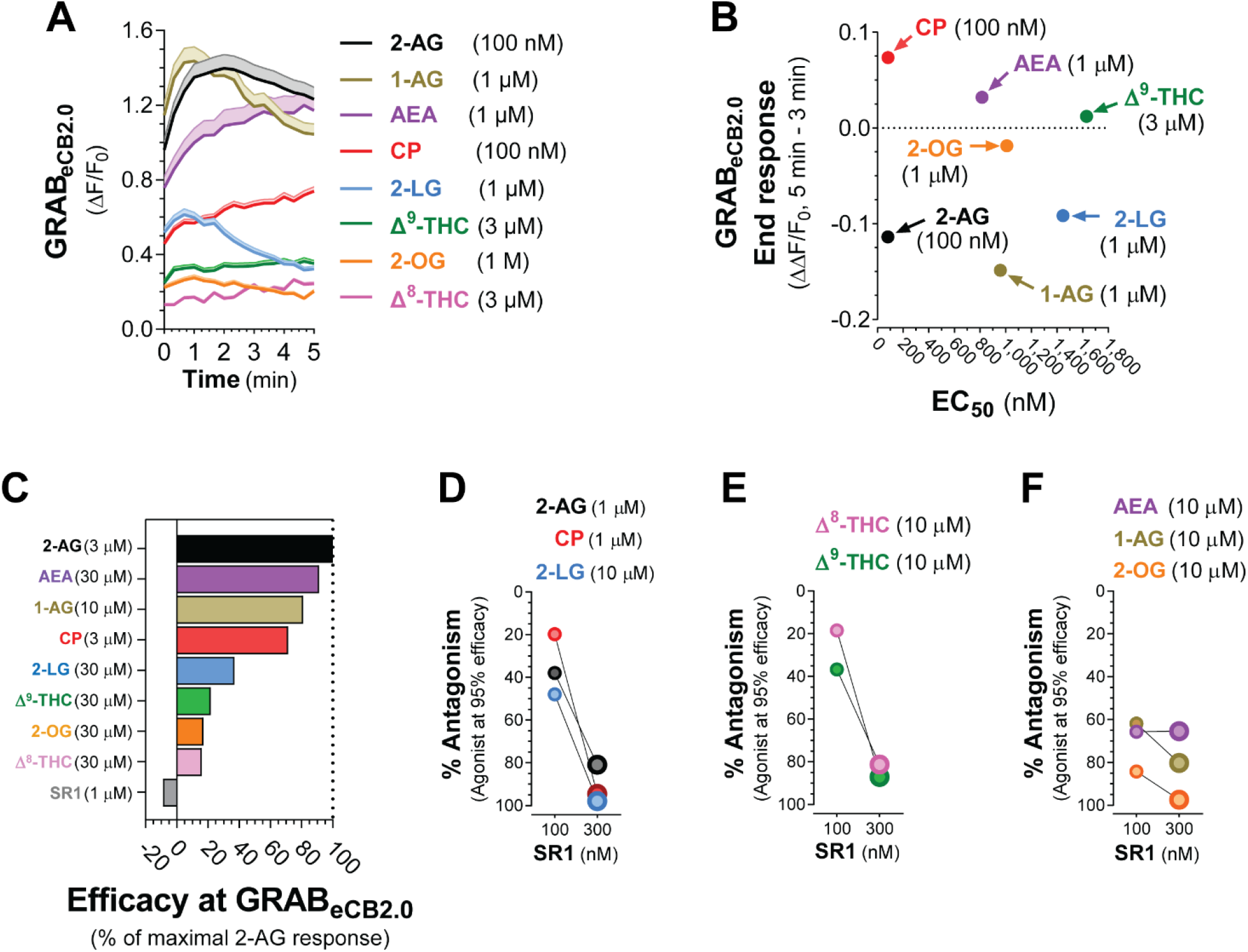
Monoacylglycerols have distinct dynamics and pharmacological properties at GRAB_eCB2.0_ compared to THC, CP, and AEA. HEK293 cells were transfected with *eCB2.0* DNA plasmid and changes in fluorescent signal measured by <u>high-throughput fluorescence assay</u>. **A]** Comparison of GRAB_eCB2.0_ activation kinetics elicited by each ligand at a concentration that approaches their respective EC_50_s. Shaded areas in time courses and error bars represent s.e.m. **B]** Comparison of the EC_50_ of each ligand and their corresponding end response at a concentration that approached their respective EC_50_ and measured between 3-5 min. Y axis positive value represent increase in fluorescent signal (i.e., AEA, CP and D^9^-THC) and negative values represent decrease in fluorescent signal (i.e., 2-AG, 1-AG, 2-LG and 2-OG). **C]** Comparison of the maximum increase in GRAB_eCB2.0_ fluorescent signal as expressed by area under the curve and as a percent of 3 μM 2-AG produced by each ligand. **D-F]** Antagonism of each response by SR1 (100 and 300 nM).

## Discussion

### Similarities and differences in GRAB_eCB2.0_ and CB_1_R pharmacology

GRAB_eCB2.0_ was developed starting from CB_1_R and screening for genetic constructs and individual randomized mutations to improve the change in fluorescent signal in response to 2-AG.^17^ The initial pharmacological characterization of GRAB_eCB2.0_ was performed in the absence of BSA in the buffer and resulted in EC_50_s values that are lower than their activities at CB_1_R: 2-AG = 3-9 μM, AEA = 0.3-0.8 μM, CP = 20 nM and THC = 2 μM).^15^ BSA is known to facilitate the solubility and interaction of CB agents with CB_1_R.^33^ We found that inclusion of BSA in the buffer improves 2-AG’s potency at GRAB_eCB2.0_ to 82 nM compared to published values, but didn’t affect the activities of AEA, THC and CP at GRAB_eCB2.0_ compared to published values. This result suggests that BSA might preferentially facilitate the solubility and interaction of select CB agents with CB_1_R and GRAB_eCB2.0_, here 2-AG.

2-AG interacts with specific amino acids within the orthosteric binding site of CB_1_R that are likely conserved in GRAB_eCB2.0_.^5;38^ By contrast, AEA, eCB analogues, phyto-CBs and artificial CBs bind to different amino acids within the orthosteric binding site that might have been mutated in GRAB_eCB2.0_ and reduce their interaction with this sensor compared to CB_1_R. Thus, our study suggests that, compared to CB_1_R, GRAB_eCB2.0_ expressed by cells in culture and mouse tissues reliably senses nanomolar changes in 2-AG levels, and micromolar amounts of AEA, eCB analogues, THC and CP.

### GRAB_eCB2.0_ pharmacology relative to abundance of eCB analogues and CB agents

Diacylglycerol lipase (**DAGL**) produces several monoacylglycerols in addition to 2-AG that exhibit significant activity at CB_1_R, such as 2-LG.^11;12^ Furthermore, monoacylglycerol lipase (**MAGL**) and α/β domain containing 6 (**ABHD6**) hydrolyze several MAGs in addition to 2-AG, including 2-LG and 2-OG.^43^ However, 2-LG is 10 less abundant than 2-AG and requires 100-fold higher concentrations than 2-AG to activate CB_1_R (EC_50_ ≈ 25 μM). Considering these values, we conclude that the activity-dependent increases in GRAB_eCB2.0_ fluorescent signal measured in cells in culture and mouse tissues that are blocked by DAGL inhibitors and increased by MAGL/ABHD6 inhibitors are much more likely to involve change in 2-AG levels than changes in 2-LG and 2-OG levels.

Δ^9^-THC increases GRAB_eCB2.0_ fluorescent signal with an EC_50_ that is 100-fold higher than its activity at CB_1_R. Intraperitoneal injections of Δ^9^-THC (5 mg/kg) in adult mice results in ≈10 ng/ml of THC in mouse brain (31.8 nM).^44;45^ Accordingly, we conclude that GRAB_eCB2.0_ expressed in mouse brain will therefore be able to sense i.p. injections of Δ^9^-THC and Δ^8^-THC and enable studying their PK profile and how this correlates with changes in behavior, or when either ligand arrives at a particular circuit or brain region in real time.

### GRAB_eCB2.0_ plateau versus decay dynamics

When applied at a concentration that approaches their EC_50_ values, CB agents increased GRAB_eCB2.0_ fluorescent signal with different kinetics: 2-AG, 1-AG, 2-LG, and 2-OG triggered rapid increases in fluorescent signal that were followed by slow decays, whereas AEA, CP, Δ^9^-THC and Δ^8^-THC triggered progressive increases in fluorescent signal that did not reach peak response within 5 min. HEK293 cells do not express MAGL, ABHD6 and fatty acid amid hydrolase (**FAAH**) but may express an unidentified hydrolase known to metabolize monoacylglycerols,^46;47^ an enzymatic activity that is likely involved in the slow decay response of 2-AG, 1-AG, 2-LG, and 2-OG. HEK293 cells also lack CYP2C9 and CYP3A4 enzymes that metabolize Δ^9^-THC and Δ^8^-THC. Thus, Δ^9^-THC and Δ^8^-THC are likely not enzymatically degraded by HEK293 cells in culture and will constantly increase GRAB_eCB2.0_ fluorescent signal.

### NAM activity of CBD at GRAB_eCB2.0_

Allosteric binding sites are distinct protein domains from orthosteric sites that bind small molecules and either increase or decrease orthosteric site-mediated changes in protein conformations and activities. Early in vitro and in vivo studies showed that CBD reduces CB_1_R signaling at concentrations well below its reported affinity (Ki) to the orthosteric agonist site of CB_1_R, providing the initial evidence that CBD acts as a NAM at this receptor.^48–50^ Accordingly, in neurons in cell culture, CBD reduces the efficacy and potency of 2-AG and THC at increasing CB_1_R signaling and inhibits eCB-mediated synaptic plasticity without influencing basal neurotransmission.^51–53^ Mutagenesis of CB_1_R indicated that several N-terminal residues of the CB_1_R, namely Cys98, Cys107, and Met1, interact with CBD when it occupies the putative allosteric site of the CB_1_R that CBD may target, and that this allosteric site overlaps with the orthosteric site which is near the second extracellular loop.^8;54^ In silico modeling of CB_1_R with an intact N-terminus revealed a potential binding pocket for NAMs of CB_1_R in close proximity its N terminus, one of the longest among class A GPCRs, and molecular docking studies suggest that binding to this site may results in a change in the 3 dimensional structure of the orthosteric binding site and thus in THC’s and 2-AG’s potencies.^54;55^ While these results suggest that CBD inhibits CB_1_R signaling by directly interacting with an allosteric binding site on this target, direct demonstration of CBD binding to CB_1_R is still needed. We found that CBD does not affect baseline GRAB_eCB2.0_ fluorescent signal but reduces 2-AG’s activity at GRAB_eCB2.0_, a result consistent with the premise that the molecular mechanism involved in mediating CBD’s allosteric modulation of CB_1_R remains functional in GRAB_eCB2.0_. Accordingly, mutagenesis optimization of CB_1_R constructs to generate GRAB_eCB2.0_ did not significantly affect its N-terminus (Supplementary Figure S1). Our results suggest that structural and mechanistic comparisons of CBD activity at CB_1_R and GRAB_eCB2.0_ might help us better understand that molecular mechanism involved in the allosteric modulation of CB_1_R. That is-how allosteric ligands produce a distinctive receptor conformation with unique signaling and therapeutic value.

### In conclusion

We show that 2-AG increases GRAB_eCB2.0_ fluorescent signal as a full agonist and with an EC_50_ similar to its activity at CB_1_R, and that GRAB_eCB2.0_ responds to AEA, 2-LG and 2-OG with EC_50_s that are lower than their EC_50_s at CB_1_R. Considering the lower amount of AEA, 2-LG and 2-OG produced by cells, our results suggest that activity-dependent increases in GRAB_eCB2.0_ fluorescent signal measured in cells in culture and mouse tissues will mainly reflect change in 2-AG levels, especially when this response is blocked by inhibitors of 2-AG production and inactivation. We also show that SR1 blocks GRAB_eCB2.0_ fluorescent signal with an IC_50_ similar to its reported potency at CB_1_R, and that this leads to levels below baseline fluorescent signal, suggesting that GRAB_eCB2.0_ exhibits basal fluorescence, and that SR1 may act as an inverse agonist and represents a useful pharmacological tool to validate GRAB_eCB2.0_ functionality when expressed by various model systems. THC and CP increase GRAB_eCB2.0_ fluorescent signal with EC_50_s lower than their activity at CB_1_R indicating that only high brain concentration of these agents will be detected in cell culture and mouse tissue model systems. CBD reduces the 2-AG-induced increase in GRAB_eCB2.0_ fluorescent signal but not its basal fluorescent signal, suggesting that the molecular mechanism of CBD allosterism present in CB_1_R is maintained in GRAB_eCB2.0_. Thus, GRAB_eCB2.0_ provides an opportunity to study how changes in THC and CBD concentration and co-activity at CB_1_R might occur in cell culture and mouse tissue model systems. Our results outline the pharmacological profile and activation dynamics of GRAB_eCB2.0_ to improve the interpretation of changes in its fluorescent signal when expressed in various model systems.

## Funding Statement

This work was supported by the National Institutes of Health (NS118130 and DA047626 to N.S., DA055448 to AE, DA033396 to MRB, and T32GM007750). We would also like to acknowledge support from the University of Washington Center of Excellence in Opioid Addiction Research/ Molecular Genetics Resource Core (P30DA048736). National Natural Science Foundation of China (31925017 and 31871087), the Beijing Municipal Science & Technology Commission (Z181100001318002 and Z181100001518004), the NIH BRAIN Initiative (1U01NS113358), the Shenzhen-Hong Kong Institute of Brain Science (NYKFKT2019013), the Science Fund for Creative Research Groups of the National Natural Science Foundation of China (81821092) and grants from the Peking-Tsinghua Center for Life Sciences and the State Key Laboratory of Membrane Biology at Peking University School of Life Sciences (to YL).

## Authorship Contribution

**Simar Singh:**Conceptualization, Methodology, Validation, Investigation, and Writing-Original Draft, and Visualization. **Dennis Sarroza:** Investigation. **Maya McGrory:** Formal analysis and software. **Anthony English:** Formal analysis and software. **Ao Dong and Yulong Li:** Resources and Writing-Review & Editing. **Larry Zweifel:** Resources and Writing-Review & Editing. **Michael R. Bruchas:** Conceptualization and Writing-Review & Editing. **Benjamin B. Land:** Conceptualization and Writing-Review & Editing. **Nephi Stella:** Conceptualization, Writing-Original Draft, Writing-Review & Editing, Visualization, Supervision, and Funding Acquisition.

## Conflict of Interest

The authors declare that they do not have any known competing financial interests or relationships that could have influenced the work in this paper.

## Notes

### Competing Interest Statement

The authors have declared no competing interest.

