## Supplementary figures and images for "Pharmacological characterization of the endocannabinoid sensor GRAB_eCB2.0_"

### Supplemental Figure 1

#### Slide 1
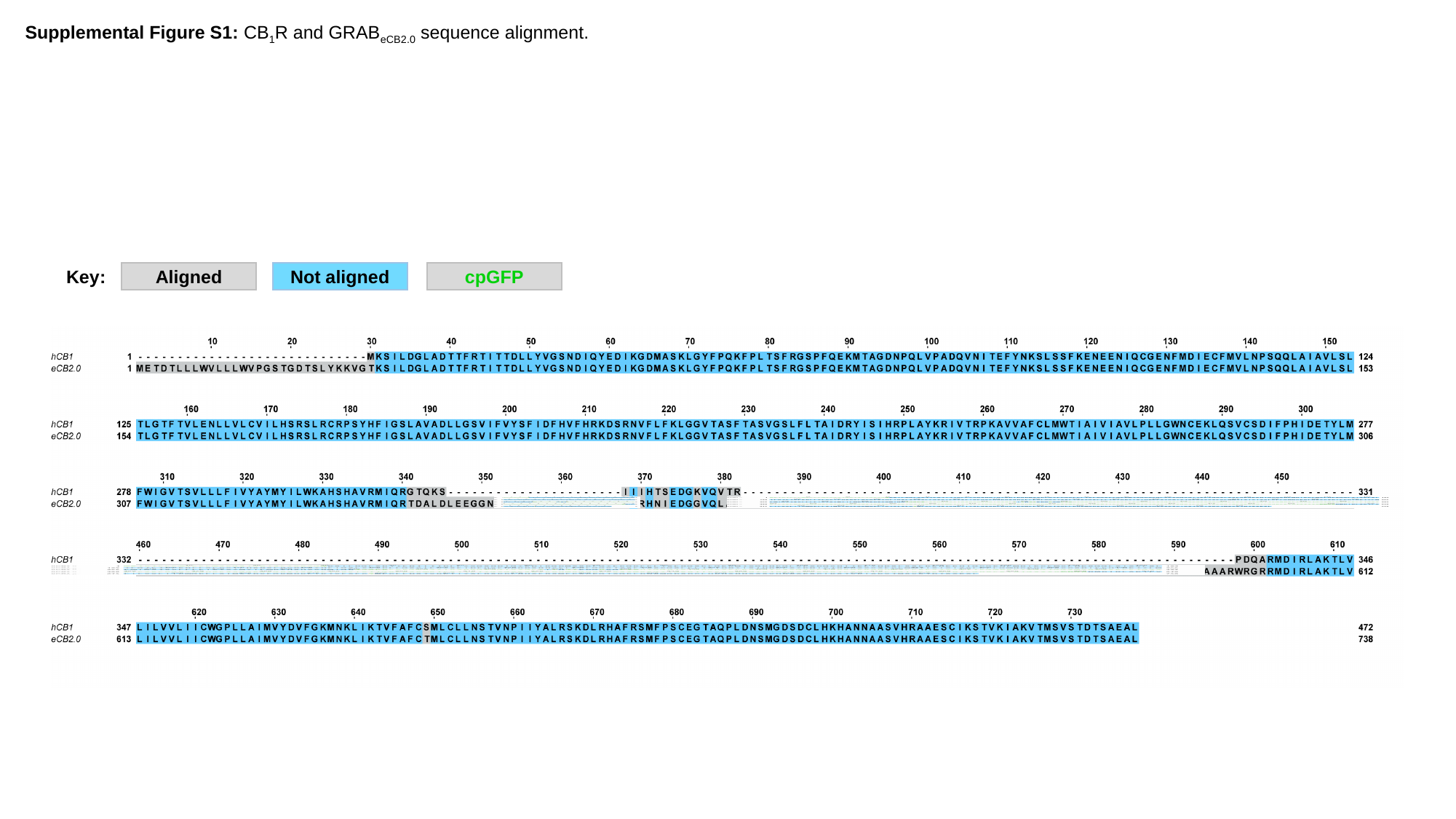

Supplemental Figure S1: CB_1_R and GRAB_eCB2.0_ sequence alignment.
Key:
Aligned
Not aligned
cpGFP
