## Supplemental Figure 4 for "Pharmacological characterization of the endocannabinoid sensor GRAB_eCB2.0_"

### Slide 1
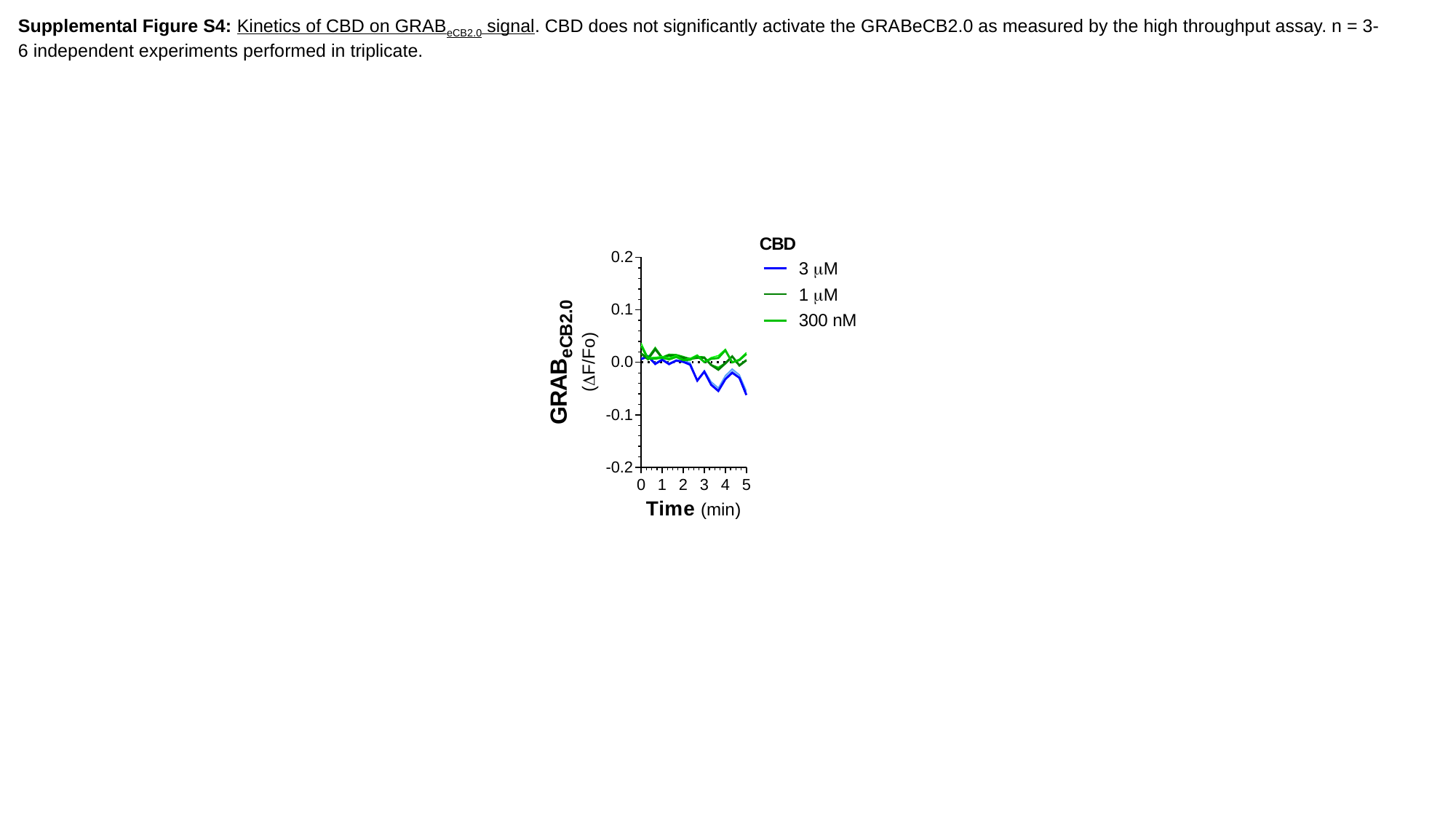

Supplemental Figure S4: Kinetics of CBD on GRAB_eCB2.0_ signal. CBD does not significantly activate the GRABeCB2.0 as measured by the high throughput assay. n = 3-6 independent experiments performed in triplicate.
