## Supplemental Figure 5 for "Pharmacological characterization of the endocannabinoid sensor GRAB_eCB2.0_"

### Slide 1
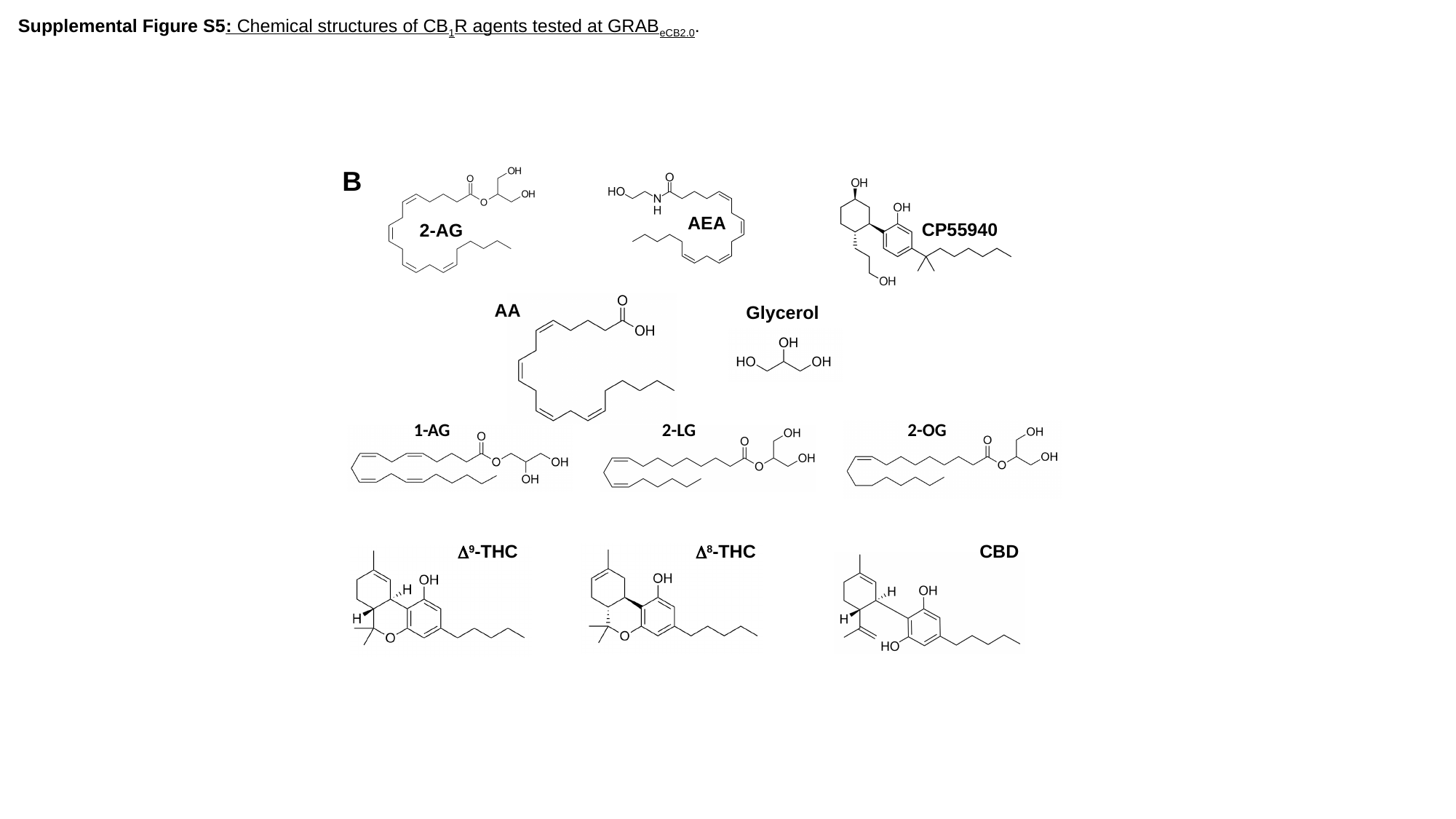

Supplemental Figure S5: Chemical structures of CB_1_R agents tested at GRAB_eCB2.0_.
B
AEA
CP55940
2-AG
HEK 293
AA
Glycerol
1-AG
2-LG
2-OG
Δ^9^-THC
Δ^8^-THC
CBD
